# Female sex hormones exacerbate retinal neurodegeneration

**DOI:** 10.1101/2024.07.11.603104

**Authors:** Ashley A. Rowe, Sofia Reyes, Mauricio J. Velasquez, Tiffany Yee, Emily R. Nettesheim, Jeffrey G. McDonald, Katherine J. Wert

## Abstract

Neurodegenerative disorders such as Alzheimer’s disease and macular degeneration represent major sources of human suffering, yet the factors influencing disease severity remain poorly understood. Sex has been implicated as one potential modifying factor. Here, we show that female sex is a risk factor for worsened outcomes in a model of retinal degeneration. Further, we show that this susceptibility is caused by the presence of female-specific circulating sex hormones. The adverse effect of female sex hormones was specific to diseased retinal neurons, and depletion of these hormones ameliorated this phenotypic effect. These findings provide novel insights into the pathogenesis of neurogenerative diseases and how sex hormones can impact the severity of disease. These findings have far-reaching implications for clinical trial design and the use of hormonal therapy in females with certain neurogenerative disorders.

## Introduction

Sex differences play critical roles in the pathogenesis of neurodegenerative diseases, such as Alzheimer’s disease, Parkinson’s disease, and age-related macular degeneration (*1-5*). Sexual differences in disease severity can be the result of many factors, including differences in anatomy, sex hormone concentrations, differentially expressed genes, and epigenetic differences between males and females (*6*). In several forms of neurodegeneration, it has been shown that women experience worsened outcomes compared to men, but the reasons for this remain poorly understood. Thus, there is an urgent need to close the gap in our understanding of these sex-related differences in order to provide better care and treatment for women with these conditions.

Retinitis Pigmentosa (RP) is a retinal neurodegeneration in which the photoreceptors, the primary light sensing neurons responsible for vision, undergo a progressive and irreversible degeneration and eventual death. As these important retinal cells die, patients experience vision loss that initiates as night blindness and progresses to full blindness. In this study, we used the well-documented mouse model of autosomal dominant RP, the most common form of this neurodegenerative disease, caused by a proline to histidine substitution at position 23 of rhodopsin (Rho P23H) (*7*) to investigate whether sex differences impact the retinal degenerative phenotype. We found that female sex was associated with more rapid neurodegeneration, and that this effect was caused by female-specific circulating sex hormones.

Current knowledge remains limited on sex hormones and whether they play a role in the mammalian retina, in either healthy or diseased states. As sex hormones are among the most widely prescribed medications for women in the United States (*8, 9*), there is a critical need to understand how such a widely prescribed medication affects chronic conditions and re-evaluate the safety of these medications for patients with certain forms of RP and other neurodegenerative disorders. Our data highlight a novel systemic hormone-dependent interaction of the female sex hormones with the pathology and progression of an inherited retinal dystrophy that has previously thought to be unaffected by biological sex.

## Results

### Female sex is associated with severity of retinal neurodegeneration

We performed electroretinography (ERG) to test the visual function of Rho P23H female and male mice. Mice underwent scotopic ERG every month through seven months of age. While both male and female mice had continued neurodegeneration over the seven months, consistent with the RP phenotype, female mice displayed a significantly faster decline in visual function beginning at two months of age. 0.01 cd.s/m^2^ scotopic recordings, indicative of rod photoreceptor function, were significantly lower in female mice compared to male littermates through seven months of age (**Fig. 1A**). 1.0 cd.s/m^2^ scotopic recordings followed a similar outcome, where female mice had significantly lower visual response in both the b- (**Fig. 1E**) and a-wave amplitudes (**Fig. 1F**). Representative traces show the loss of phototransduction in female Rho P23H mice compared to male littermates (**Fig. 1B-D, G-I)**. We next analyzed the structural changes in the retina to assess if the sexual dimorphic nature was solely a functional loss in the photoreceptor neurons or whether it reflected loss of cell survival. We found that female Rho P23H mice had a significantly thinner outer nuclear layer (ONL), comprising the photoreceptor nuclei, than male counterparts at seven months of age (**Fig. 1J, K**).

**Figure 1.**
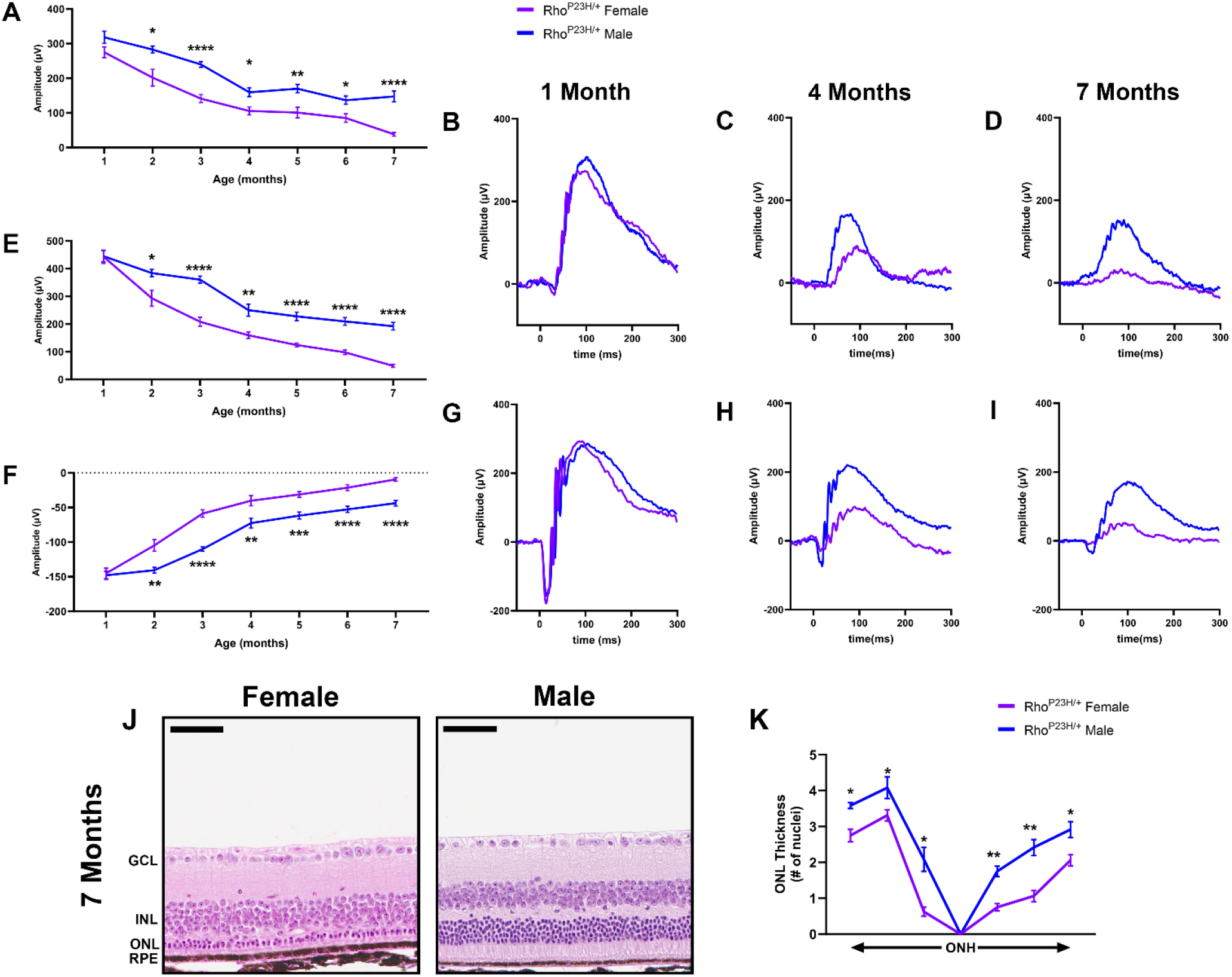
Rho P23H mice display a sexually dimorphic loss of photoreceptor function and survival. **(A)** 0.01 cd•s/m^2^ scotopic ERG b-wave values in male (blue) and female (purple) Rho P23H mice from one to seven months of age. **(B-D)** Representative 0.01 cd•s/m^2^ scotopic ERG traces at one, four, and seven months of age. **(E)** 1.0 cd•s/m^2^ scotopic ERG b- and **(F)** a-wave amplitudes. **(G-I)** Representative 1.0 cd•s/m^2^ scotopic ERG traces at one, four, and seven months of age. Error bars = SEM. *, p<0.05. **, p<0.01. ***, p<0.001. ****, p<0.0001. Statistical analysis performed by multiple unpaired *t*-test with a Holm-Šídák Correction with α=0.05. N=16 eyes per group. **(J)** Representative H&E-stained histology from seven-month Rho P23H female and male mouse retinas. **(K)** Quantification of outer nuclear layer (ONL) thickness at six different regions spanning from the optic nerve head (ONH). GCL, ganglion cell layer; INL, inner nuclear layer; RPE, retinal pigmented epithelium. Statistics performed via multiple unpaired *t*-tests and a Holm-Šídák multiple comparisons test. *, p-value<0.05; **, p-value<0.01. Scale bar = 50 µM. N ≥ 3 mice per group.

### Removal of gonadal sex hormones depletes hormone levels within the mammalian retina

Our observed sex difference in visual function began at two months of age, after mice reach sexual maturity and after the onset of disease in this model system (approximately postnatal day 18) (*10-12*). To assess the effect of the steroidal sex hormones on retinal neurodegeneration, we surgically removed the ovaries (bilateral ovariectomy; OVX) and testis (orchiectomy; OCX) from Rho P23H female and male mice at sexual maturity (six weeks of age; **Fig. 2A**). This gonadectomy approach results in removal of the primary sex hormone-producing organs but leaves the remainder of the hypothalamic-pituitary-gonadal axis intact. We employed mass spectrometric analysis of mouse serum and neural retinal samples to assess hormone concentrations with and without surgical manipulation. At eight weeks of age, two weeks post-surgery, serum exhibited lowered testosterone and progesterone after gonadectomy (**Fig. 2B**). The retina also displayed detectable and significant changes in hormone concentrations similar to that found in the serum samples (**Fig. 2C**).

**Figure 2.**
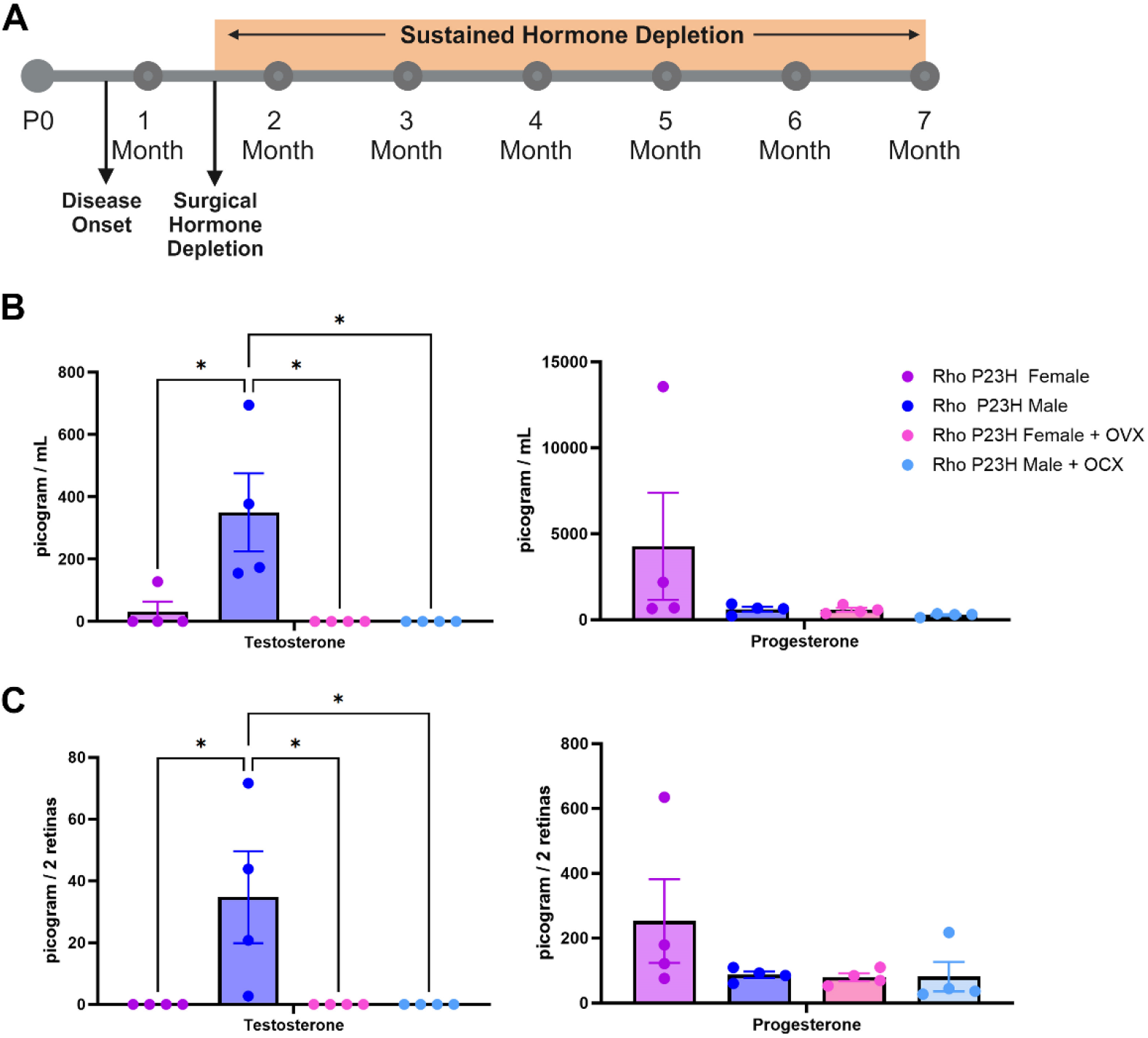
Surgical gonadectomy depletes steroidal sex hormones in the serum and retina. **(A)** Timeline depicting surgical procedure at six weeks of age, after disease onset, followed by sustained hormone depletion through the duration of the study. P0, post-natal day zero. **(B)** Mass spectrometry analysis of serum and **(C)** retina testosterone and progesterone in Rho P23H females (purple), males (blue), females with bilateral ovariectomy (OVX; pink), and males with orchiectomy (OCX; light blue), two-weeks post-surgery. *, *p* < 0.05. Statistics performed via One-way ANOVA with Tukey’s multiple comparison’s test. N = 4 mice per group. Part of figure made with Biorender.com (agreement #QH271R0NYW).

### Female sex hormones worsen retinal neurodegeneration

We assessed the effect of systemic hormone depletion on Rho P23H RP visual function by ERG. Analysis of the 0.01 cd.s/m^2^ scotopic setting revealed that there were no appreciable differences between intact males and males who received OCX. However, females that received OVX had a significant increase in visual function compared to intact females, which was not significantly different from males (**Fig. 3A-C**). 1.0 cd.s/m^2^ scotopic ERG showed similar results, where females with OVX had significantly higher ERG values for both the b- (**Fig. 3D, F, G**) and a-wave amplitudes (**Fig. 3E-G**), matching that of male littermates.

**Figure 3.**
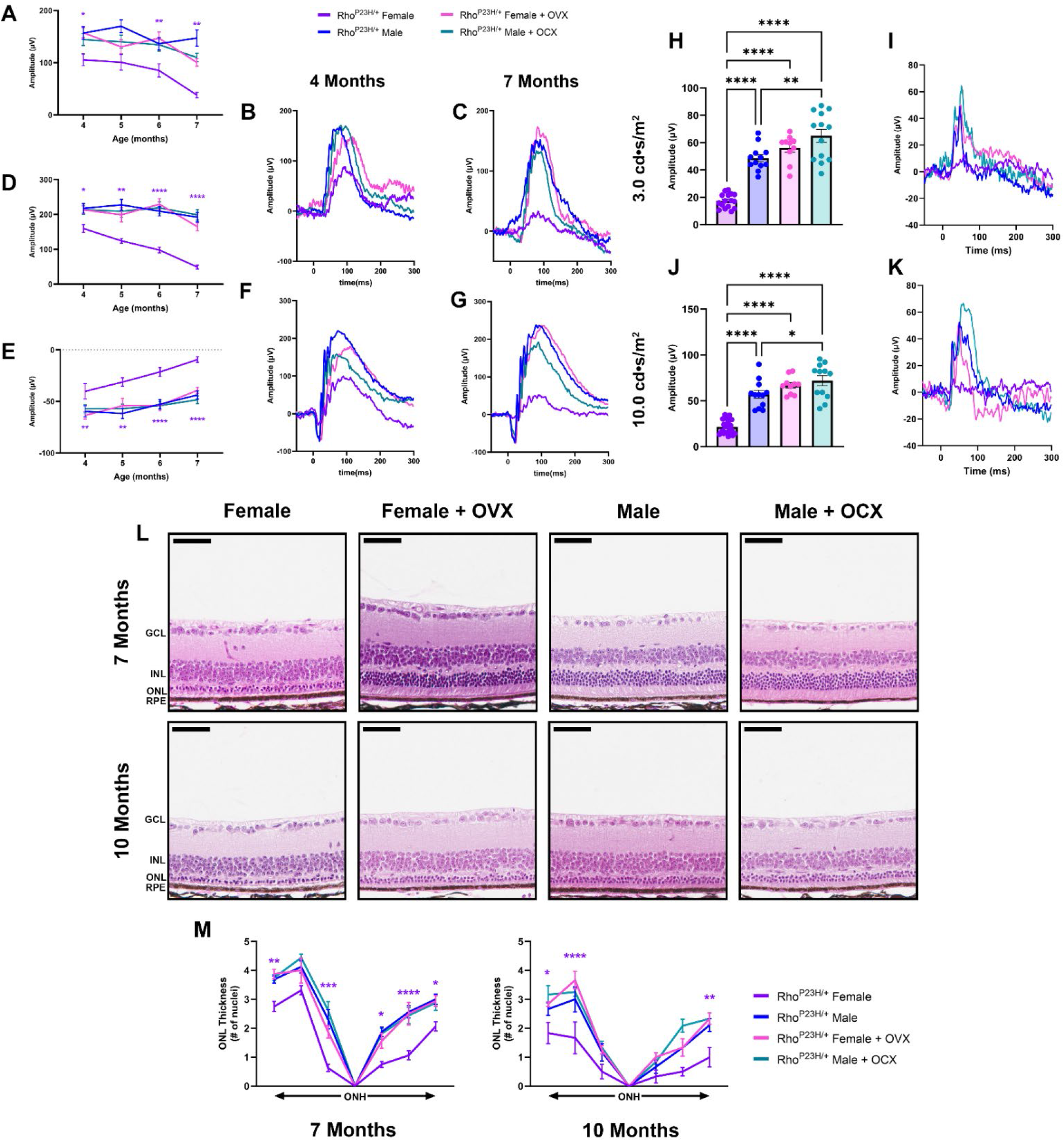
Ovariectomy ameliorates female photoreceptor neurodegeneration. **(A)** 0.01 cd•s/m^2^ scotopic b-wave amplitudes from four to seven months of age in Rho P23H females (purple), males (blue), females with OVX (pink), and males with OCX (light blue). **(B-C)** Representative 0.01 cd•s/m^2^ scotopic traces. **(E)** 1.0 cd•s/m^2^ scotopic b- and **(F)** a-wave amplitudes from four to seven months of age, and **(F-G)** representative traces. Statistics performed via Two-way ANOVA with Tukey’s multiple comparison’s test. Statistical results are shown for females compared to females + OVX and males compared to males + OCX. N≥16 eyes. **(H)** 3.0 cd•s/m^2^ and **(J)** 10.0 cd•s/m^2^ photopic amplitudes at seven months of age, **(I, K)** with respective representative traces. Statistics analyzed by One-way ANOVA with Tukey’s multiple comparison’s test. N≥10 eyes. Error bars=SEM. **(L)** H&E-stained histology from seven and ten-month-old Rho P23H female and male mice with and without OVX or OCX, respectively. **(M)** Quantification of ONL thickness spanning from the ONH at both seven and ten months of age. Statistics performed via Two-way ANOVA with Tukey’s multiple comparison’s test. Statistical results are shown for females compared to females + OVX. N≥3 mice per group. Scale bar=50 µM. *, p<0.05; **, p<0.01; ***, p<0.001; ****, p<0.0001.

In RP, as the rod photoreceptor neurons die, the cone photoreceptors experience secondary death, leading to a loss of central vision in later stages of retinal degeneration. We assessed photopic signal at late-stage retinal degeneration in the Rho P23H mice to investigate cone neuron-specific function. Both 3.0 cd.s/m^2^ (**Fig. 3H, I**) and 10.0 cd.s/m^2^ (**Fig. 3J, K**) revealed that female mice had significantly reduced photopic signal, while females with OVX retained a higher photopic signal similar to that of male photopic amplitudes. We did note that male mice with OCX had a significantly higher photopic reading compared to intact males at both light intensity settings (**Fig. 3H-K**). We next assessed the late structural changes in the retina and found that male mice who received OCX had no difference in ONL thickness compared to intact males. Female mice who received OVX had a significantly thicker ONL than intact females at both seven and ten months of age, matching the photoreceptor survival found in male mice and congruent with the ERG results (**Fig. 3L, M**).

### Female sex hormones impact degenerating but not healthy retinal neurons

To determine whether the steroidal sex hormones impact the retinal neurons outside of a disease context, we performed OVX on C57BL/6J wild-type females and assessed visual function by ERG over time compared to intact males and females. We found no difference between female mice that had received OVX and intact females or males through seven months of age for 0.01 cd.s/m^2^ scotopic and 1.0 cd.s/m^2^ scotopic b- and a-wave amplitudes (**Fig. 4A-E**). We also analyzed retinal histology from these mice at seven months of age. No difference was observed for ONL thickness in any of the six regions analyzed throughout the retina (**Fig. 4F, G**).

**Figure 4.**
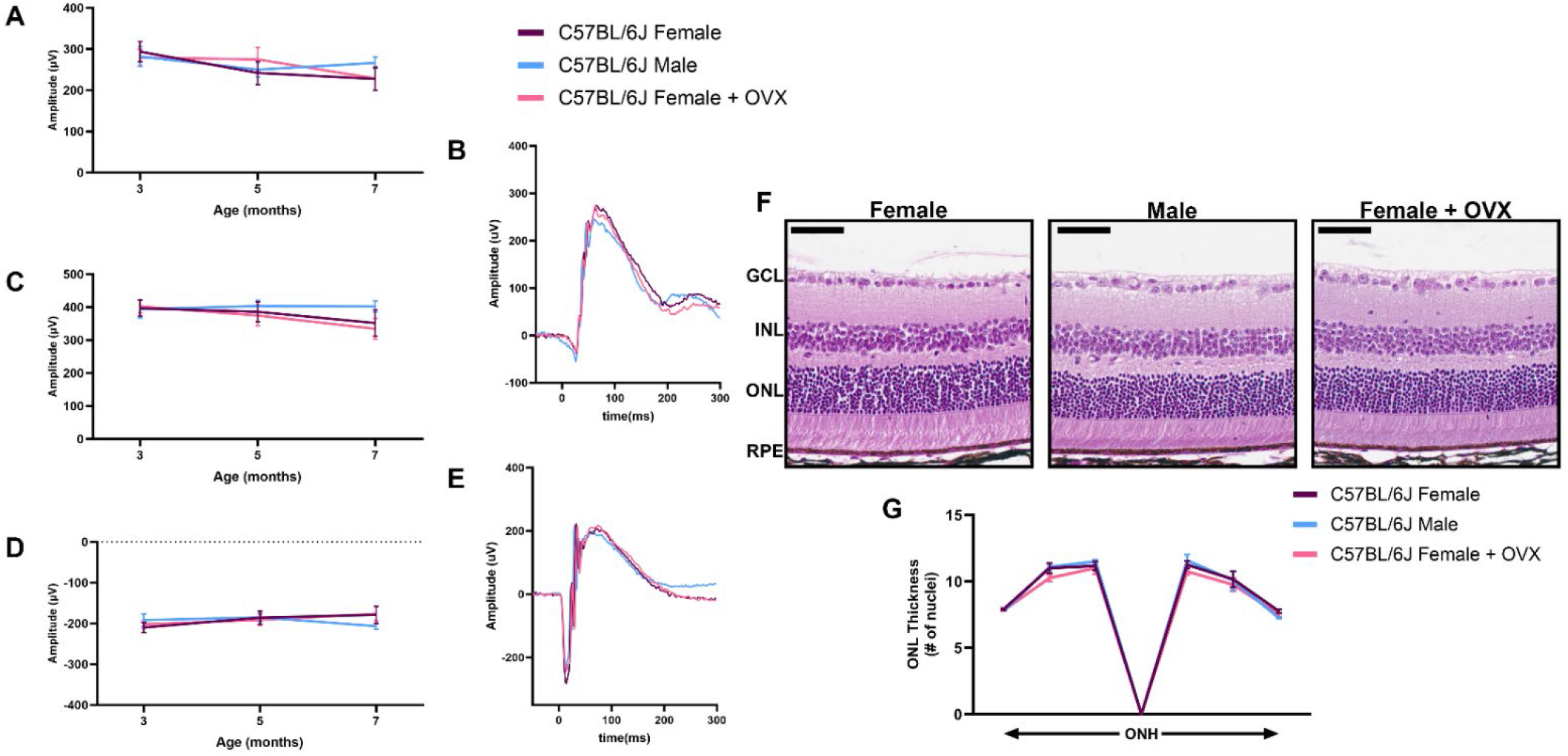
Sex hormone depletion does not affect healthy retinal neurons. **(A)** 0.01 cd•s/m^2^ scotopic ERG b-wave amplitudes, **(C)** 1.0 cd•s/m^2^ scotopic ERG b- and **(D)** a-wave amplitudes from male (blue), female (purple), and females with bilateral ovariectomy (OVX; pink) C57BL/6J mice at three, five and seven months of age. Representative ERG traces for the **(B)** 0.01 cd•s/m^2^ and **(E)** 1.0 cd•s/m^2^ scotopic ERGs settings for all three groups at seven months of age. Statistics analyzed via Two-way ANOVA (α=0.05) with Tukey’s multiple comparison’s test. N = 10 eyes per group. **(F)** H&E-stained histology of retinas from seven-month-old C57BL/6J females, males, and females + OVX. **(G)** Quantification of the outer nuclear layer (ONL) thickness at six different regions in the retina spanning out from the optic nerve head (ONH) for all groups. GCL, ganglion cell layer; INL, inner nuclear layer; RPE, retinal pigmented epithelium. Statistics performed via Two-way ANOVA with Tukey’s multiple comparison’s test. Scale bar = 50 µM. N = 3 eyes per group.

## Discussion

Investigation into mechanisms that have profound effects on underlying neurodegeneration in an incurable disease such as RP is of great clinical interest as the field seeks to find ways to slow neuronal loss and preserve function. In this study, we investigated and defined the sexually dimorphic vision loss between male and female mice with Rho P23H knock-in mutations that mimic the most prevalent form of autosomal dominant RP. We determined the contribution of sex-specific hormones on the manifestation of sex differences in Rho P23H RP, as hormonal interactions present opportunities for therapeutic intervention. Through hormone depletion studies *in vivo*, we found that depletion of the female sex hormones resulted in a near total amelioration of the functional and structural sex difference in our RP model of neurodegeneration.

Sex hormones carry out biological function in various cells of the body by binding to hormone receptors either on the cell surface or intracellularly (*13-22*). Prior studies have shown mRNA and protein for these receptors being present in retinal tissue (*23*), but their exact function within retinal neurons is unknown. Evidence that hormone signaling may play a role in development of retinal tissue as well as other neural tissue, might explain why some these proteins are present during development, but their role in physiology and disease in the mature retina and brain remains elusive. Studies in retinal degenerative disease report beneficial effects of hormone therapy (*23-29*). One group found that topical administration of progesterone to the eye resulted in an alleviation of oxidative stress and reduction in apoptosis in the *rds* mouse model of RP (*29*). In this study, a sexually dimorphic phenotype was not reported, and the study was mainly conducted at post-natal day 21, before the onset of sexual maturity. Another study reported a neuroprotective effect of 17-β-Estradiol on the retinas of mice undergoing NMDA retinal toxicity, but they assessed 6–8-week-old male mice (*23*). Our data suggests that an interaction of hormone signaling with genetic mutations causal for neurodegenerative disease reduces function and neuronal cell survival in females. We determined that depletion of female sex hormones does not impact retinal neuron function or survival in healthy retina, and therefore the hormones interact within the cell stress response or cell death mechanisms underlying the rhodopsin genetic mutation. Future studies into hormone interconversions and receptors that mediate their signaling in the retina will be needed to further define how this signaling impacts the photoreceptor neurons.

The majority of human clinical studies investigate patients carrying several forms of neurodegenerative disease, such as RP, grouped together in longitudinal analysis, but individual mutation analysis is not typically considered. When looking into specific genetic mutations, one study published that mutations in *Pde6β* caused retinas from female mice to degenerate on a faster timeline (*30*), and a recent study supports our observations on sex differences, albeit with no link to mechanism (*31*). As other retinal degenerative diseases such as age-related macular degeneration and diabetic retinopathy have noted sex differences, our data argues that biological sex as a variable should be rigorously tested in preclinical models to avoid biasing of resultant data. Overall, our data highlight a novel systemic hormone dependent interaction of the female sex hormones with the pathology and progression of an inherited retinal dystrophy that has previously thought to be unaffected by biological sex (**Fig. 5**). These important findings highlight the need to advocate for sex stratification in natural history and clinical trial studies for patients with RP and other neurodegenerative diseases and mark the importance for record keeping of hormonal medications for these patients.

**Figure 5.**
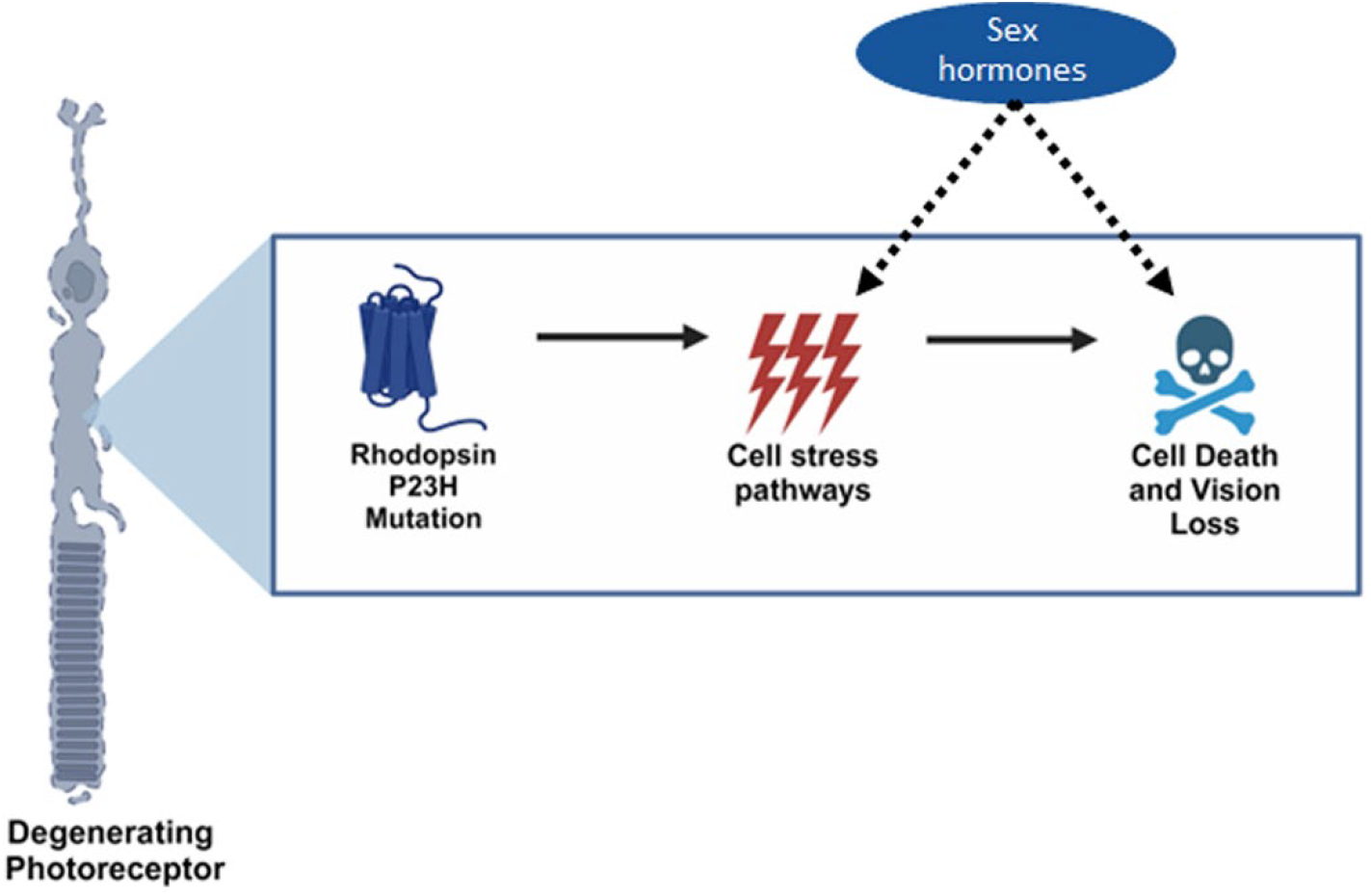
Sex hormones impact cellular pathways underlying the rhodopsin P23H mutation in photoreceptor neurons. Graphical schematic depicting our findings that female systemic sex hormones interact within pathways found in diseased, but not healthy, retinal neurons to exacerbate neurodegeneration in a sexually dimorphic manner. Part of figure made with BioRender.com (agreement #XT271R0LO4).

## Materials and Methods

### Mouse lines and husbandry

All experiments were performed in accordance with the Annual Association for Research in Vision and Ophthalmology (ARVO) Statement for the Use of Animals in Ophthalmic and Visual Research and were all approved by the Animal Care and Use Committee at UT Southwestern Medical Center (UTSW). Rho P23H (RRID: IMSR_JAX:017628) mice were obtained, carrying a point mutation that mimics the human autosomal dominant form of Rho P23H retinitis pigmentosa as previously described. Rho P23H homozygous mice were bred with C57BL/6J mice (Jackson Laboratory, Bar Harbor ME; RRID: MSR_JAX:000664) to obtain the mice used in this study. C57BL/6J mice were used as wild-type controls. Additionally, heterozygous Rho P23H breeding was used, and littermates were analyzed for the sexual dimorphism in disease, which occurred only in mice carrying the P23H mutation in rhodopsin. All mice were maintained in approved animal facilities at UTSW and were kept on a normal light–dark cycle (12/12 h). Food and water were available *ad libitum* throughout the experiment. Experiments were conducted monthly, with a month being measured as a cycle of 28 days.

### Electroretinography (ERG)

Mice were dark-adapted for at least 12 h, manipulations were conducted under dim red-light illumination, and recordings were made using the Celeris ERG system by Diagnosys LLC (Lowell, MA, USA) following previous methods (*37, 38*). Briefly, pupils were dilated using topical 2.5% phenylephrine hydrochloride (Akorn Inc.) and 1% tropicamide (Sandoz). Mice were anesthetized by intraperitoneal injection of 0.1 mL/10 g body weight of anesthesia (1 mL ketamine 100 mg/mL and 0.1 mL xylazine 20 mg/mL in 8.9 mL 1x PBS). Body temperature was maintained at 37°C during the testing. The mouse’s eyes were covered in a 0.3% hypromellulose gel (GENTEAL, Alcon). Scotopic retinal responses were recorded at 0.01 and 1.0 cd•s/m^2^ white light intensity settings. A minimum of eight measurements were recorded and averaged for each light setting. Photopic ERG was recorded at seven months of age, immediately following the scotopic ERG using the Celeris system. First, mouse eyes were exposed to 10 min of white light, followed by flashes at two white light settings: 3.0 and 10.0 cd•s/m^2^, each one averaging at least fifteen sweeps.

### Gonadectomies

Ovariectomy and orchiectomy were performed as previously described (*39*). Briefly, mice were shaved, and subcutaneous pain medications of meloxicam (5 mg/kg) and buprenorphine SR (1.0 mg/kg) were delivered. Anesthesia was maintained using 2% isoflurane via precision vaporizer. Eye surfaces were protected using artificial tear ointment and body temperature was maintained at 37°C throughout the procedure. Orchiectomy was performed via a single midline incision in the ventral scrotal sac where both testis were removed. Ovariectomy was performed via two dorsal-lateral incisions into the abdominal wall where the ovaries were isolated and severed by the distal uterine horn, resulting in removal of the oviduct and ovary. In both cases, internal layers were closed using 4-0 absorbable suture material and skin wounds were closed with wound clips. Confirmation of ovariectomy success was monitored via estrus cycle cessation observed via vaginal cytology and visual identification of estrus phases (*40*).

### Retinal histology and quantification

Mice were sacrificed following institutional guidelines. Enucleation was performed by proptosing the eye and placing a curved pair of forceps behind the eye, near the optic nerve, and gently pulling outward, releasing the eye and a portion of the optic nerve as previously described (*37, 38*). Briefly, eyes were fixed at room temperature and embedded in paraffin, sectioned, and stained with H&E by Excalibur Pathology Inc. Each section contained upper and lower retina as well as the posterior pole. Retinal sections were imaged using light microscopy (Hamamatsu NanoZoomer S60, UTSW Whole Brain Microscopy Facility RR:SCR_017949). Quantification of photoreceptor nuclei was conducted on several sections of the eye that contained the optic nerve, as follows: the distance between the optic nerve and the ciliary body was divided into three, approximately equal, regions on each side of the eye. The number of nuclei in four columns was counted within each single region for the outer nuclear layer (ONL) of the retina. These counts were then used to determine the average thickness of the ONL for each individual animal and within each region of the retina, spanning from the ciliary body to the optic nerve head and back out to the ciliary body.

### Retina dissection and serum collection

Prior to mass spectrometry analysis, blood was collected by submandibular bleeding using a 5mm lancet. Whole blood was collected into a 1.5 mL uncoated centrifuge tube and allowed to coagulate at room temperature for 30 minutes to an hour. Next, the sample was spun in a 4°C centrifuge for 10 min at 1000xg. The serum supernatant was pipetted off the top and frozen at - 80°C until ready for processing. After blood collection, the mice were euthanized via cervical dislocation prior to dissection of the retinas. A razor blade was used to slice a midline incision into the cornea. Next, a curved pair of forceps was used to remove the lens, followed by removal of the neural retina, which was placed in a power bead tubes (Qiagen) and flash frozen in liquid nitrogen. Retinas were stored at -80°C until ready for processing.

### Mass Spectrometry

Retinas: Retinas were placed in bead beater tubes (PowerBead Tubes, Ceramic 2.8mm, Qiagen, Germany) with 200uL of water and 20uL of stable isotope labeled steroid cocktail and homogenized using 5.50ms^-1^, 10s speed time (Bead Ruptor 24, Omni, Kennesaw, GA). Then 400uL of cold methanol was added to each sample, vortexed for 30s, and then let stand for 5 min. Samples were centrifuged at 4415 × G for 5 min. Steroids were isolated from the sample using solid phase extraction (Evolute Express ABN, (30 mg/1mL), Biotage, Charlotte, NC). Columns were sequentially conditioned with 1mL of acetonitrile, 1mL of methanol, and 0.5mL water. All SPE work was carried out with a positive pressure manifold (Biotage, Charlotte, NC) operated at 6 psi. Samples were loaded onto the conditioned columns and washed with 0.5mL water and then 0.5mL of 30% methanol, both of which were discarded. Steroids were eluted with two 0.5mL aliquots of acetonitrile into a 2mL polypropylene plate (2mL Square 96-Well Plate with 100uL Tapered Reservoir, Analytical Sales, Flanders, NJ). Samples were evaporated under a gentle stream of nitrogen at 40 °C and reconstituted in 100uL of 50% methanol.

#### Serum

An 100uL aliquot of serum was added to a 2mL microcentrifuge tube along with 300uL of water and 20uL of stable isotope labeled steroid cocktail. The mixture was vortexed for 5 s and then 200uL of ZnSO_4_ was added, and it was vortexed again for 10 s. A 400uL aliquot of cold methanol was added, the samples were vortexed for 30 s, then let stand for 5 minutes and then centrifuged at 4415 × G for 5 min. Steroids were isolated using the same SPE procedure described above for the retina samples. Samples were reconstituted with 200uL of 50% methanol.

Steroids were quantitatively measured using high performance liquid chromatography – mass spectrometry as previously described (*41*).

### Statistical Analysis

Data are reported as mean ± SEM unless otherwise noted. GraphPad Prism Software (version 10.0) was used to generate graphs and perform statistical analysis. Data comparing three or more groups were analyzed using one-way ANOVA followed by Tukey’s multiple comparison’s test. Data comparing two groups over multiple time points were analyzed using multiple two-tailed *t*-tests with the Holm-Šidák’s method to correct for multiple comparisons. Data with two independent variables were analyzed using two-way ANOVA followed by Tukey’s multiple comparison’s test. For all applicable statistical tests, α was set to 0.05. Normal distribution and homogeneity of variance was determined by graphical analysis. See individual methods sections, results and figure legends for specific testing methods. No mice were excluded from the analysis. A *p* value of less than 0.05 was considered significant and measurements were done

## Acknowledgements

We thank the members of the Department of Ophthalmology, the Sexual Dimorphism and Sex Hormone Signaling in Disease Research Interest Group, and the Genetics, Development and Disease graduate program at UT Southwestern Medical Center for their valuable advice and guidance. We also thank Dr. David Birch and other members of the Retina Foundation of the Southwest (Dallas, TX) for their input related to RP patients. We thank the Whole Brain Microscopy Facility for their assistance in imaging the mammalian retina.

## Author contributions

Conceptualization: AAR, KJW

Methodology: AAR, KJW

Investigation: AAR, MJV, TY, ERN

Visualization: AAR, SR, KJW

Validation: AAR, MJV, JGM, KJW

Funding acquisition: JGM, KJW

Supervision: KJW

Writing: AAR, KJW

## Data and materials availability

All data are available in the main text or the supplementary materials.

